# SARS-CoV-2 neutralising antibodies in Dogs and Cats in the United Kingdom

**DOI:** 10.1101/2021.06.23.449594

**Authors:** Shirley L. Smith, Enyia. R. Anderson, Cintia Cansado-Utrilla, Tessa Prince, Sean Farrell, Bethaney Brant, Stephen Smyth, Peter-John M. Noble, Gina L. Pinchbeck, Nikki Marshall, Larry Roberts, Grant L. Hughes, Alan D. Radford, Edward I. Patterson

**Author notes:** These authors contributed equally to this work (Shirley L. Smith, Enyia. R. Anderson, Alan D. Radford, Edward I. Patterson). Corresponding author: Edward I. Patterson, Department of Biological Sciences, Brock University, St. Catharines, ON L2S 3A1, Canada.

## Abstract

Companion animals are susceptible to SARS-CoV-2 infection and sporadic cases of pet infections have occurred in the United Kingdom. Here we present the first large-scale serological survey of SARS-CoV-2 neutralising antibodies in dogs and cats in the UK. Results are reported for 688 sera (454 canine, 234 feline) collected by a large veterinary diagnostic laboratory for routine haematology during three time periods; pre-COVID-19 (January 2020), during the first wave of UK human infections (April-May 2020) and during the second wave of UK human infections (September 2020-February 2021). Both pre-COVID-19 sera and those from the first wave tested negative. However, in sera collected during the second wave, 1.4% (n=4) of dogs and 2.2% (n=2) cats tested positive for neutralising antibodies. The low numbers of animals testing positive suggests pet animals are unlikely to be a major reservoir for human infection in the UK. However, continued surveillance of in-contact susceptible animals should be performed as part of ongoing population health surveillance initiatives.

## Introduction

Severe acute respiratory syndrome coronavirus-2 (SARS-CoV-2) emerged in Wuhan, China at the end of 2019 [1] and rapidly spread around the world. The main route of transmission remains human-to-human. However, there is evidence that the virus can infect animals [2] and it is important that we remain vigilant of such infections; particularly in companion animals with whom humans often have close contact.

Although initially there were only sporadic cases of infection in cats and dogs [3–5], there are now numerous reports of infection detected by RT-PCR or virus isolation [6–10], including in the UK [11]. Evidence of infection of cats and dogs has also been provided by the detection of anti-SARS-CoV-2 antibodies in several studies; from Italy, France, Germany, Croatia and China [12–17]. Experimental infections have shown that cats and, to a lesser extent, dogs are susceptible to SARS-CoV-2 and that cats can transmit the virus to other cats [18–20]. Infections in companion animals appear to have occurred as a result of human-to-animal transmission; however, the reported transmission of SARS-CoV-2 from farmed mink to in-contact humans, cats and dogs [21, 22] and the detection of the virus in stray dogs and cats [23, 24], suggest it is important to continue surveillance in companion animals. Here we conducted a survey of SARS-CoV-2 neutralising antibodies in cats and dogs attending UK veterinary practices.

## Methods

### Samples

Canine and feline sera used in this study were obtained from the UK Virtual Biobank, which uses health data from commercial diagnostic laboratories participating in the Small Animal Veterinary Surveillance Network (SAVSNET) to target left over diagnostic samples in the same laboratories for enhanced phenotypic and genomic analyses [25]. All samples were residual sera remaining after routine diagnostic testing and were sent by the contributing laboratory based on convenience within the following parameters: samples were requested from UK cats and dogs collected over two time periods; March and April 2020 (early pandemic) for both cats and dogs, then September 2020 to February 2021 for dogs, and January 2021 for cats (late pandemic). Serum samples collected from the same laboratory in early January 2020 were also tested as pre-COVID-19 controls. All samples were linked to electronic health data for that sample (species, breed, sex, postcode of the submitting veterinary practice, date received by the diagnostic laboratory) held in the SAVSNET database, using a unique anonymised identifier. Data on SARS-CoV-2 exposure or symptoms was not available. Ethical approval to collect electronic health data (SAVSNET) and physical samples from participating laboratories (National Virtual Biobank) was granted by the Research Ethics Committee at the University of Liverpool (RETH000964).

### Neutralising antibody detection in serum samples

Serum samples were screened for SARS-CoV-2 neutralising antibodies using the plaque reduction neutralisation test (PRNT) as previously described [15], with the SARS-CoV-2/human/Liverpool/REMRQ0001/2020 isolate cultured in Vero E6 cells [26]. Briefly, sera were heat inactivated at 56°C for 30 mins and stored at −20°C until use. DMEM containing 2% FBS was used to dilute sera ten-fold followed by serial two-fold dilution. SARS-CoV-2 at 800 plaque forming units (PFU)/ml was added to diluted sera and incubated at 37°C for 1 h. The virus/serum mixture was then inoculated onto Vero E6 cells, incubated at 37°C for 1 h, and overlaid as in standard plaque assays [27]. Cells were incubated for 48 h at 37°C and 5% CO_2_, fixed with 10% formalin and stained with 0.05% crystal violet solution. PRNT_80_ was determined by the highest dilution with 80% reduction in plaques compared to the control. Samples with detectable neutralising antibody titre were repeated as technical replicates for confirmation. Where titres differed between technical replicates, the lowest dilution was reported.

## Results

A total of 732 samples were received from the diagnostic laboratory and tested for SARS-CoV-2 neutralising antibodies. Linking of data to the samples found that 22 samples were duplicates (duplicate samples gave the same result in each replicate and are therefore reported as one sample). Seven samples were from animals with non-UK postcodes, two samples did not have species data, two samples were received as dogs but were actually from cats and were collected outside the two time periods of cat sample collection and eleven samples were missing postcodes; these samples were excluded. Results are therefore reported for 688 sera (454 canine, 234 feline) of which 558 (372 dogs, 186 cats) were collected during the SARS-CoV-2 pandemic and 130 (82 dogs, 48 cats) were collected from animals before the first confirmed human case in the UK (21^st^ January 2020 [28]) - pre-COVID-19 samples; these samples were distributed across the UK (Figure 1). Of the dog sera collected during the pandemic, 0/85 (0%) collected in March/April 2020 and 4/287 (1.4%) collected September 2020-February 2021 tested positive for neutralising antibodies with titres ranging from 1:20 to 1:80. In cats, 0/96 (0%) sera collected in March/April 2020 tested positive for neutralising antibodies and 2/90 (2.2%) collected in January 2021 tested positive with titres of 1:40 and 1:80. Pre-COVID-19 sera from both dogs (n=82) and cats (n=48) tested negative for neutralising antibodies. Positive samples in dogs were collected in November 2020 (n=1), January 2021 (n=2) and February 2021 (n=1) and were collected in Kent, Buckinghamshire, Worcestershire and Yorkshire, respectively (Figure 1). The two positive cats were collected in January 2021; one in Birmingham and the other in London (Figure 1).

**Figure 1:**
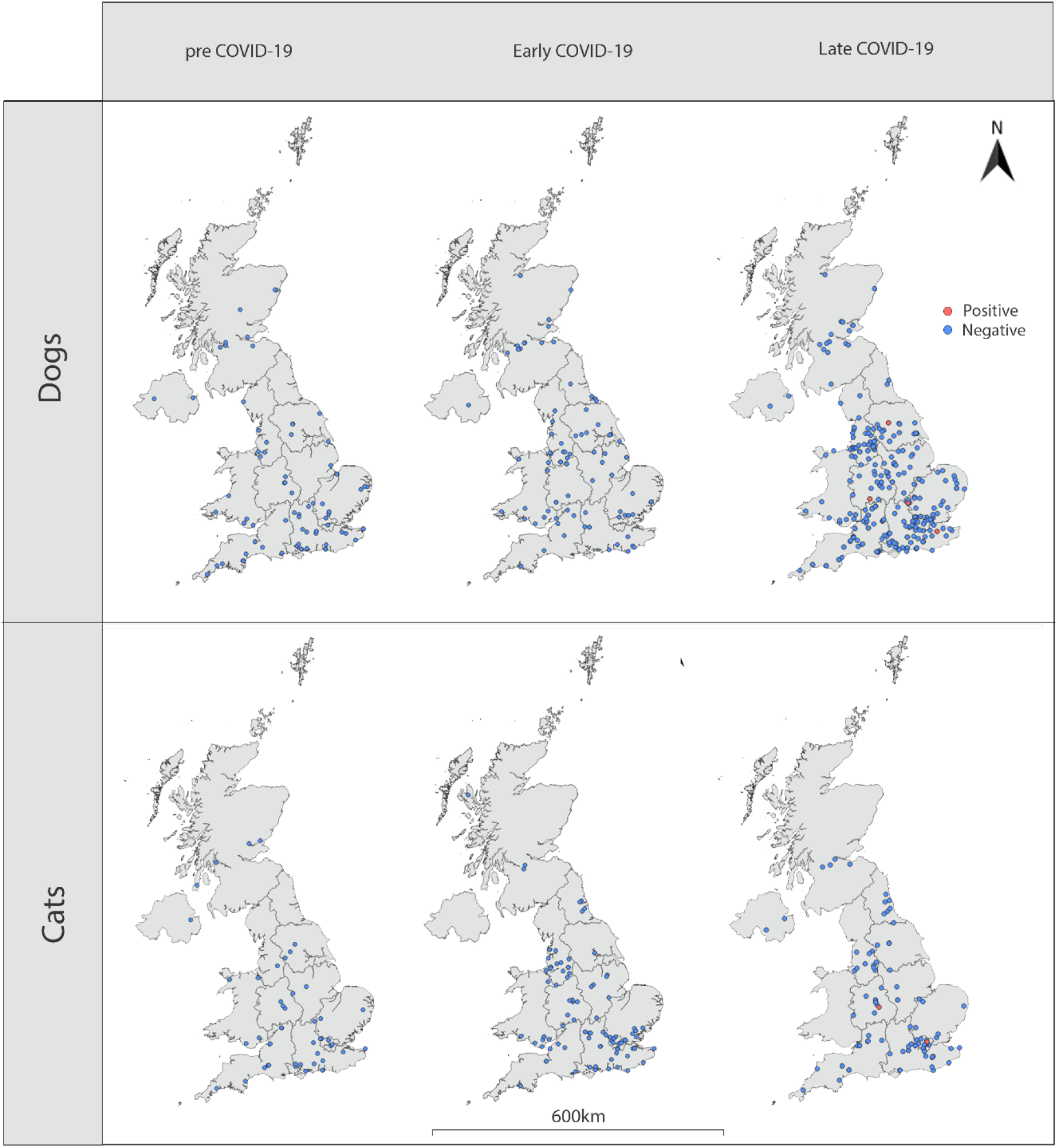
Schematic map showing the location of samples for which testing of SARS-CoV-2 neutralising antibodies is reported. Red dots indicate samples that were positive for SARS-CoV-2 neutralising antibodies using PRNT_80_. Blue dots indicate samples that were negative.

## Discussion

SARS-CoV-2 emerged in humans in China late in 2019, rapidly spreading across the world. Studies of companion animals from several countries have shown that they too can be infected with the virus. In the UK, there are sporadic reports of infection in cats and dogs [11, 29], however, there has been no large scale test of infection. Here we show that a small proportion of UK dogs and cats sampled at a time of active human transmission tested positive for SARS-CoV-2 neutralising antibodies.

Sera from two time points during the pandemic were analysed. Sera collected early in the pandemic, during March and April 2020, from both cats and dogs were negative for neutralising antibodies. Previous studies using European samples have shown a low level of infection, highest in Italy, where 3.3% (15/451) of dog sera and 5.8% (11/191) cat sera collected between March and May 2020 had measurable neutralising antibody titres [15]. These samples were purposefully collected from regions of Italy with a high prevalence of infection in humans, in some cases from households known to contain recently diagnosed human cases. Our results in contrast, are more consistent with a survey from a similar population of cats in Germany, that found 0/221 samples collected in April and May of 2020 to be positive for anti-SARS-CoV-2 antibodies using ELISA [13], and with a survey in the Netherlands in April-May 2020, that found 0.4% of cats and 0.2% dogs to be seropositive [30]. Lack of positive samples from this time period in the UK (April-May2020) likely reflects the selection criteria of the animals assayed (undergoing routine haematological testing and not selected based on location), and the relatively low rate of human disease at the time compared to Italy.

In sera collected later in the pandemic, 4/287 (1.4%) dogs and 2/90 (2.2%) cats tested positive. Positive dog samples were collected in November 2020 and January and February of 2021. Positive cats were collected in January 2021. This is again broadly in line with a recent German survey conducted from September 2020 to February 2021, showing a seroprevalence of 1.36%, that the authors concluded corresponded with the rise of reported cases in the human population, and was suggestive of ongoing transmission from owners to their cats [14].

Cats and dogs can be infected with other coronaviruses, leading to the possibility that SARS-CoV-2 neutralising antibodies in cats and dogs may result from previous infection with a different virus. We and others have previously demonstrated a lack of cross-reactivity between SARS-CoV-2 and samples containing antibodies to feline coronavirus (FCoV), canine enteric coronavirus (CeCoV) and canine respiratory coronavirus (CRCoV) [13, 15, 16]; all of which are endemic in UK cats and dogs [31–33]. Here we also tested samples from UK cats and dogs collected before the human index case in the UK (21^st^ January 2020 [28]). All pre-COVID-19 samples were negative for SARS-CoV-2 neutralising antibodies. Similar results have been reported for both cats and dogs by others [30], suggesting that antibodies produced following infection by cat and dog coronaviruses do not cross react with SARS-CoV-2.

Here we made use of samples collected from a commercial diagnostic laboratory contributing data to a voluntary national surveillance scheme (SAVSNET) to efficiently test for evidence of prior SARS-CoV-2 infection in UK cats and dogs. The major limitations of such a system are the relatively sparse data available for each sample such that individual animals, that are not identifiable, may have been sampled twice or have come from the same household. In addition, such samples lack detailed information on the health of the animals and whether they were from a COVID-19-positive household. However, acquiring such samples from the UK Virtual Biobank, offers a responsive resource for studying national patterns of disease in UK pets [25].

We report here the detection of SARS-CoV-2 neutralising antibodies during the second wave of human infections in the UK. Other groups have previously reported that cats and dogs can become infected, likely through their interactions with humans. Although animal-to-animal transmission has been reported, for example on mink farms and in experimental infections [18–20, 22, 34], the small numbers of companion animals testing positive in the field suggest that pets are not currently acting as a significant reservoir for infection, and that the pandemic will be controlled by measures largely focussed on minimising human-to-human transmission. However, studies like that presented here strongly argue for continued surveillance of in-contact, susceptible animal species, which will help determine whether in the future, more targeted control measures are needed for pet animals, particularly in regions that are gaining control of infection in their human populations.

## References

1. Wu F, Zhao S, Yu B, et al. A new coronavirus associated with human respiratory disease in China. Nature 2020; 579:265–9.

2. Prince T, Smith, S.L., Radford, A.D., Solomon, T. Hughes, G.L. and Patterson, E. I. SARS-CoV-2 Infections in Animals: Reservoirs for Reverse Zoonosis and Models for Study. Viruses 2021; 13.

3. Garigliany M, Van Laere AS, Clercx C, et al. SARS-CoV-2 Natural Transmission from Human to Cat, Belgium, March 2020. Emerg Infect Dis 2020; 26:3069–71.

4. Newman A, Smith D, Ghai RR, et al. First Reported Cases of SARS-CoV-2 Infection in Companion Animals - New York, March-April 2020. MMWR Morb Mortal Wkly Rep 2020; 69:710–3.

5. Sit THC, Brackman CJ, Ip SM, et al. Infection of dogs with SARS-CoV-2. Nature 2020.

6. Barrs VR, Peiris M, Tam KWS, et al. SARS-CoV-2 in Quarantined Domestic Cats from COVID-19 Households or Close Contacts, Hong Kong, China. Emerg Infect Dis 2020; 26:3071–4.

7. Decaro N, Vaccari G, Lorusso A, et al. Possible Human-to-Dog Transmission of SARS-CoV-2, Italy, 2020. Emerg Infect Dis 2021; 27.

8. Hamer SA, Pauvolid-Correa A, Zecca IB, et al. SARS-CoV-2 Infections and Viral Isolations among Serially Tested Cats and Dogs in Households with Infected Owners in Texas, USA. Viruses 2021; 13.

9. Ruiz-Arrondo I, Portillo A, Palomar AM, et al. Detection of SARS-CoV-2 in pets living with COVID-19 owners diagnosed during the COVID-19 lockdown in Spain: A case of an asymptomatic cat with SARS-CoV-2 in Europe. Transbound Emerg Dis 2020.

10. Sailleau C, Dumarest M, Vanhomwegen J, et al. First detection and genome sequencing of SARS-CoV-2 in an infected cat in France. Transbound Emerg Dis 2020; 67:2324–8.

11. Hosie MJ, Epifano I, Herder V, et al. Detection of SARS-CoV-2 in respiratory samples from cats in the UK associated with human-to-cat transmission. Vet Rec 2021; 188:e247.

12. Fritz M, Rosolen B, Krafft E, et al. High prevalence of SARS-CoV-2 antibodies in pets from COVID-19+ households. One Health 2021; 11:100192.

13. Michelitsch A, Hoffmann D, Wernike K, Beer M. Occurrence of Antibodies against SARS-CoV-2 in the Domestic Cat Population of Germany. Vaccines (Basel) 2020; 8.

14. Michelitsch A, Schon J, Hoffmann D, Beer M, Wernike K. The Second Wave of SARS-CoV-2 Circulation-Antibody Detection in the Domestic Cat Population in Germany. Viruses 2021; 13.

15. Patterson EI, Elia G, Grassi A, et al. Evidence of exposure to SARS-CoV-2 in cats and dogs from households in Italy. Nat Commun 2020; 11:6231.

16. Stevanovic V, Vilibic-Cavlek, T., Tabain, I., Benvin, I., Kovac, S., Hruskar, Z., Mauric, M., Milasincic, L., Antolasic, L., Skrinjaric, A., Staresina, V. and Barbic, L. Seroprevalence of SARS-CoV-2 infection among pet animals in Croatia and potential public health impact. Transbound Emerg Dis 2020.

17. Zhang Q, Zhang H, Gao J, et al. A serological survey of SARS-CoV-2 in cat in Wuhan. Emerg Microbes Infect 2020; 9:2013–9.

18. Bosco-Lauth AM, Hartwig AE, Porter SM, et al. Experimental infection of domestic dogs and cats with SARS-CoV-2: Pathogenesis, transmission, and response to reexposure in cats. Proc Natl Acad Sci U S A 2020; 117:26382–8.

19. Halfmann PJ, Hatta M, Chiba S, et al. Transmission of SARS-CoV-2 in Domestic Cats. N Engl J Med 2020.

20. Shi J, Wen Z, Zhong G, et al. Susceptibility of ferrets, cats, dogs, and other domesticated animals to SARS-coronavirus 2. Science 2020; 368:1016–20.

21. Oude Munnink BB, Sikkema RS, Nieuwenhuijse DF, et al. Transmission of SARS-CoV-2 on mink farms between humans and mink and back to humans. Science 2021; 371:172–7.

22. van Aart AE VF, Fischer EAJ, Broens EM, Egberink H, Zhao S, Engelsma M, Hakze-van der Honing RW, Harders F, de Rooij MMT, Radstake C, Meijer PA, Oude Munnink BB, de Rond J, Sikkema RS, van der Spek AN, Spierenburg M, Wolters WJ, Molenaar RJ, Koopmans MPG, van der Poel WHM, Stegeman A, Smit LAM. SARS‐CoV‐2 infection in cats and dogs in infected mink farms. Transboundary and Emerging Diseases 2021.

23. Dias HG, Resck MEB, Caldas GC, et al. Neutralizing antibodies for SARS-CoV-2 in stray animals from Rio de Janeiro, Brazil. PLoS One 2021; 16:e0248578.

24. Villanueva-Saz S, Giner J, Tobajas AP, et al. Serological evidence of SARS-CoV-2 and co-infections in stray cats in Spain. Transbound Emerg Dis 2021.

25. Smith SL, Afonso MM, Roberts L, Noble PM, Pinchbeck GL, Radford AD. A virtual biobank for companion animals: A parvovirus pilot study. Vet Rec 2021:e556.

26. Patterson EI, Prince, T., Anderson, E. I., Casas-Sanchez, A., Smith, S. L., Cansado-Utrilla, C., Turtle, L. and Hughes, G. L. Methods of inactivation of SARS-CoV-2 for downstream biological assays. 2020.

27. Rossi SL, Russell-Lodrigue KE, Killeen SZ, et al. IRES-Containing VEEV Vaccine Protects Cynomolgus Macaques from IE Venezuelan Equine Encephalitis Virus Aerosol Challenge. PLoS Negl Trop Dis 2015; 9:e0003797.

28. Lillie PJ, Samson A, Li A, et al. Novel coronavirus disease (Covid-19): The first two patients in the UK with person to person transmission. J Infect 2020; 80:578–606.

29. Ferasin L, Fritz, M., Ferasin, H., Becquart, P., Legros, V. and Leroy, E. M. Myocarditis in naturally infected pets with the British variant of COVID-19. BioRxiv Preprint 2021.

30. Zhao S, Schuurman N, Li W, et al. Serologic Screening of Severe Acute Respiratory Syndrome Coronavirus 2 Infection in Cats and Dogs during First Coronavirus Disease Wave, the Netherlands. Emerg Infect Dis 2021; 27:1362–70.

31. Addie DD, Jarrett JO. Feline coronavirus antibodies in cats. Vet Rec 1992; 131:202–3.

32. Priestnall SL, Brownlie J, Dubovi EJ, Erles K. Serological prevalence of canine respiratory coronavirus. Vet Microbiol 2006; 115:43–53.

33. Stavisky J, Pinchbeck GL, German AJ, et al. Prevalence of canine enteric coronavirus in a cross-sectional survey of dogs presenting at veterinary practices. Vet Microbiol 2010; 140:18–24.

34. Oreshkova N, Molenaar RJ, Vreman S, et al. SARS-CoV-2 infection in farmed minks, the Netherlands, April and May 2020. Euro Surveill 2020; 25.

